# Multi-modal multi-resolution atlas of the human neonatal cerebral cortex based on microstructural similarity

**DOI:** 10.1101/2022.12.14.520508

**Authors:** Mingyang Li, Xinyi Xu, Zuozhen Cao, Ruike Chen, Ruoke Zhao, Zhiyong Zhao, Xixi Dang, Kenichi Oishi, Dan Wu

## Abstract

The neonatal period is a critical window for the development of the human brain and may hold implications for the long-term development of cognition and disorders. Multi-modal connectome studies have revealed many important findings underlying the adult brain but related studies were rare in the early human brain. One potential challenge is the lack of an appropriate and unbiased parcellation that combines structural and functional information in this population. Using 348 multi-modal MRI datasets from the developing human connectome project, we found that the information fused from the structural, diffusion, and functional MRI was relatively stable across MRI features and showed high reproducibility at the group level. Therefore, we generated automated multi-resolution parcellations (300 - 500 parcels) based on the similarity across multi-modal features using a gradient-based parcellation algorithm. In addition, to acquire a parcellation with high interpretability, we provided a manually delineated parcellation (210 parcels), which was approximately symmetric, and the adjacent areas around each boundary were statistically different in terms of the integrated similarity metric and at least one kind of original features. Overall, the present study provided multi-resolution and neonate-specific parcellations of the cerebral cortex based on multi-modal MRI properties, which may facilitate future studies of the human connectome in the early development period.

## Introduction

It is widely accepted that the cerebral cortex can be divided into several subregions, which show distinct characteristics with the neighboring regions in functional or structural properties (Eickhoff et al., 2018; Glasser et al., 2016; Toga et al., 2006; Van Essen et al., 2019). A precise parcellation serves as the basis to understand the neural mechanism underlying cognitive functions and provides a standardized reference of brain areas (Glasser et al., 2016; Tzourio-Mazoyer et al., 2002). So far, several parcellations are established in human adults, created using different neurobiological features including cytoarchitecture (Amunts et al., 2020; Scholtens et al., 2015; Zilles and Amunts, 2010), sulcus/gyrus (Desikan et al., 2006; Manera et al., 2020; Tzourio-Mazoyer et al., 2002) white matter connections (Mars et al., 2011; Oishi et al., 2008; Zhang et al., 2014) and resting-state functional connections (Cohen et al., 2008; Gordon et al., 2016; Schaefer et al., 2018). Those parcellations contain areas ranging from tens to hundreds per hemisphere, enabling researchers to choose an appropriate parcellation according to their research purposes. However, limited parcellations exist for human neonates, although the neonatal stage is known to be a critical period for the development of the brain structure and function in the human brain (Eyre et al., 2021; Fenchel et al., 2020). This is an unmet need in the surge of developmental neuroscience.

Since directly warping the adults’ parcellation into neonatal space is problematic, previous studies had created a few neonate-specific parcellations based on the structural or functional features of the neonatal (Adamson et al., 2020; Alexander et al., 2017; Feng et al., 2019; Gousias et al., 2013, 2012; Oishi et al., 2019, 2011; Shi et al., 2018, 2011) see table 1 for a summary), for studying the developing connectome of the human brain. Generally, those parcellations could be classified into two types. The first type of parcellations was based on macroanatomy (sulcus or white matter bundles), and the boundaries in these atlases have clear anatomical definitions so that the results are easy to interpret and compare across different studies (Adamson et al., 2020; Alexander et al., 2017; Feng et al., 2019; Gousias et al., 2013, 2012; Oishi et al., 2011; Shi et al., 2011). However, those parcellations commonly only have tens of areas in a single hemisphere, which were too coarse for refined spatial analysis, e.g. the middle frontal or middle temporal lobes are defined as one region in UNC AAL or M-CRIB atlas (Shi et al., 2011). The other type of parcellations were based on functional connectivity (FC-based parcellation; Shi et al., 2018; Wang et al., 2021), which is particularly useful for the analysis of functional connectome. But the prior assumptions about functional homogeneity involved in the parcellation might lead to biased results when applied to a structural network (Caspers et al., 2013). Moreover, FC-based parcellation commonly has hundreds of areas in a single hemisphere, and some of those areas were asymmetric and lack relevant anatomical information, resulting in low interpretability for analysis of individual areas.

**Table 1.**
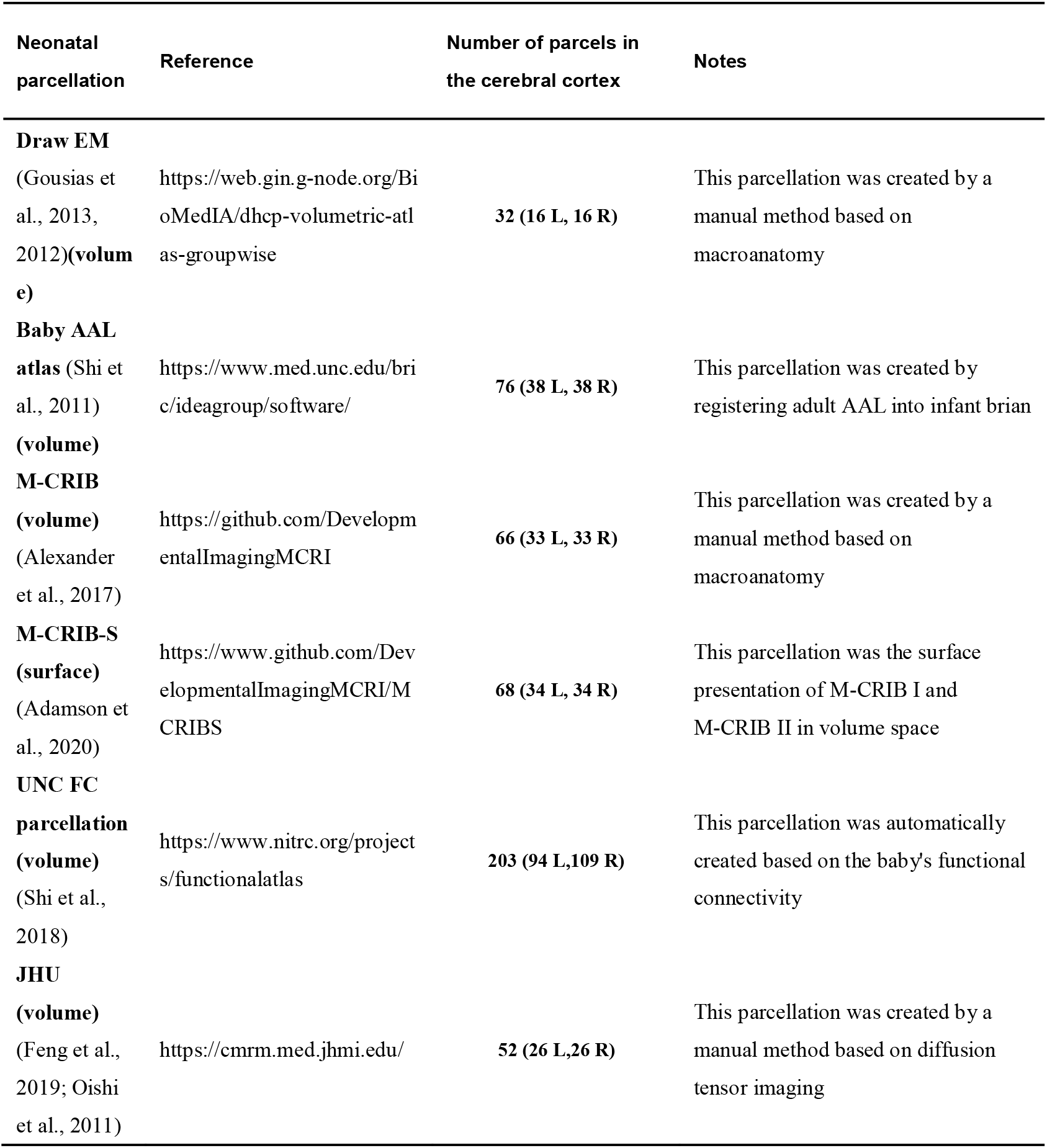
Summary of the previously published neonate-specific parcellations.

Structure-function coupling using multi-modal MRI data has been used to describe the relationship between structural and functional connectivity (Suárez et al., 2020). Previous studies revealed that the coupling between white matter and resting-state functional connectivity was a fundamental property of the brain network system, exhibited distinct spatiotemporal properties, and was related to many cognitive abilities (Baum et al., 2020; Mars et al., 2018; Vázquez-Rodríguez et al., 2019). Inspired by the intrinsic coupling and to address the aforementioned limitations in the neonatal brain atlas, we aimed to generate a comprehensive parcellation of the human neonatal cortex based on the similarity of the combined structural and functional signatures from multi-modal MRI data. The proposed atlas is expected to provide complementary and confirmatory information from multiple MRI features to identify boundaries between different areas, which would facilitate both functional and structural connectomics and also integrated structure-function analysis due to its unbiased nature (Caspers et al., 2013).

To obtain the multi-modal microstructural similarity, we used structural, diffusion, and resting-state functional MRI (r-fMRI) images of 348 neonates from the developing human connectome project (dHCP; http://www.developingconnectome.org/). We estimated the architectural (e.g. cortical thickness), microstructural (e.g. T1w/T2w myelination, tensor-based and NODDI-based measurements), and microenvironmental (ALFF and fALFF) measurements for each vertex and calculated the similarity between different vertices based on those local measurements. We used a gradient-based method (Gordon et al., 2016) to generate the final parcellations at different resolutions. One inevitable limitation of the automated parcellations is that the parcels are commonly unsymmetrical and some areas had low interpretability. Therefore, we also provided a manual parcellation based on the gradient map to ensure approximate symmetry with clear anatomical definitions.

## Methods

### Participants

A total of 783 neonatal datasets from dHCP (www.developingconnectome.org/) were collected. We excluded the subjects who 1) had high radiology scores (above 2, punctate lesions or worse) as reviewed by perinatal neuroradiologists (*n* = 218, including 2 datasets without scores); 2) were sedated during the scan (*n* = 5); 3) were scanned earlier than 37 weeks of postmenstrual age (PMA) (*n* = 91); 4) missed any of three MRI modalities including structural, diffusion and resting-state functional imaging (*n* = 160); 5) could not pass the quality control of dHCP preprocessing pipelines (*n* = 9); 6) failed the cortical registration pipeline (*n* = 4); or 7) were preterm-born neonates (birth time < 37w, *n* = 52). Finally, 348 datasets (no longitudinal data included) were selected for the present study with PMA at the scan of 41.1 ± 1.7 weeks. The dHCP study was approved by the UK Health Research Authority (14/LO/1169) and written consent was obtained from the parents or legal guardians of the study subjects.

### Data Acquisition

Multi-modal MRI scans were performed at the Evelina Newborn Imaging Centre, Evelina London Children’s Hospital on a 3-T Philips scanner with a dedicated neonatal imaging system during natural sleep. T1-weighted (T1w), T2-weighted (T2-w), resting-state functional (r-fMRI), and diffusion MRI (dMRI) images were collected within a single scan session for each neonate over 63 minutes. The detailed protocols could be found on dHCP website (https://biomedia.github.io/dHCP-release-notes/acquire.html). Briefly, the scan parameters were as follows. 1) T1w images were acquired at resolution = 0.8 × 0.8 × 1.6 mm, repetition time (TR) = 4795 ms, echo time (TE) = 8.7 ms, Inversion time (TI) = 1740 ms, SENSE factor of 2.27 (axial) and 2.66 (sagittal) and filed of view (FOV) = 145 × 122 × 100 mm. 2) T2w images were obtained at resolution = 0.8 × 0.8 × 1.6 mm, TR = 12000 ms, TE = 156 ms, SENSE factor of 2.11 (axial) and 2.58 (sagittal) and FOV = 145 × 145 × 108 mm. 3) r-fMRI were collected in 15.05 minutes with an optimized protocol for neonates (Bozek et al., 2018; Fitzgibbon et al., 2020), using multiband factor = 9, TR = 392 ms, TE = 38 ms, 2300 volumes, flip angle = 34° and spatial resolution = 2.15 mm isotropic. 4) dMRI was acquired through a spherically optimized set of directions on 4 shells (20 non-diffusion-weighted, 64 directions at b=400 s/mm^2^, 88 directions at b=1000 s/mm^2^, and 128 directions at b=2000 s/mm^2^), multiband factor = 4; TR = 3800ms; TE = 90ms; SENSE factor = 1.2; spatial resolution = 1.5 × 1.5 × 3 mm with1.5 mm overlap.

### Data Preprocessing

We collected the minimally preprocessed image data from dHCP database and the detailed preprocessing as described in the previous paper (Li et al., 2022) Briefly, for the anatomical data, the dHCP pipeline included super-resolution reconstruction (Kuklisova-Murgasova et al., 2012), registration (from T1w to T2w), bias correction, brain extraction, segmentation (Makropoulos et al., 2014), surface extraction (Schuh et al., 2017), and surface registration (Robinson et al., 2018). We used the cortical metrics obtained from dHCP, including the cortical thickness (CT) and myelination (CM) in individual brains and the corresponding transformation files from individual to the dHCP 40-weeks surface template (Bozek et al., 2018).

The functional data from dHCP included the r-fMRI data in individual spaces and the motion parameters. We further processed these data by following steps using custom codes and the DPABI toolbox (Yan et al., 2016) in MATLAB (v2018a) as described in our previous study (Li et al., 2022). 1) To decrease the influence of head motion on the data, we selected a continuous subset (1600 volumes, around 70%) of the total volume with the lowest head motion for each neonate. 2) Registration of the selected r-fMRI subset to individual T2w space. 3) Linear detrending. 4) Regression of nuisance covariates, which contained 24 head motion parameters, and the signals of white matter, cerebrospinal fluid and global brain. 5) Calculation of amplitude of low-frequency fluctuations (ALFF), fractional ALFF (fALFF) using DPABI toolbox; 6) The resulting volumes were further projected into the individual cortical surface (midthickness surface) using the ribbon-constrained procedure available in Connectome Workbench (v1.5, https://github.com/Washington-University/workbench) and then registered to the dHCP symmetric 40-week template (Robinson et al., 2018).

The dMRI data from dHCP were already preprocessed with the diffusion EDDY pipeline (Bastiani et al., 2019). Those images were further processed with the following steps: 1) registering the dMRI images to individual T2w images using FLIRT (Jenkinson et al., 2002); 2) denoising and removing Gibbs ringing artifacts with MRtrix3 tools (https://www.mrtrix.org/); 3) fitting with a diffusion tensor model to obtain axial diffusivity (AD), mean diffusivity (MD), fractional anisotropy (FA) and radial diffusivity (RD) using MRtrix3; 4) fitting with NODDI model (Zhang et al., 2012) to obtain the neurite orientation dispersion (ODI) and neurite density (or intra-cellular volume fraction, ICV) using the NODDI Matlab toolbox (http://mig.cs.ucl.ac.uk/index.php?n=Tutorial.NODDImatlab); and 5) the resulting measures were projected onto the individual cortical surfaces and then registered to the dHCP symmetric 40-week template (Bozek et al., 2018).

### Group average multi-modal local similarity map

The flowchart of generating the border density map was illustrated in Fig. 1. For each subject, there were 10 surface maps reflecting 10 different local measures from three MRI modalities, including CT, CM from T1w and T2w images; AD, FA, MD, RD of the tensor model and ICV, ODI of the NODDI model from dMRI; and ALFF, fALFF from r-fMRI. Those maps were averaged vertex-wise across subjects followed by slight smoothing with a 2mm full-width half maximum (FWHM) window, respectively. Since some of the different features shared similar spatial patterns across the whole brain (the absolute value of paired correlation among the used properties ranged from 0.01 to 0.99, median value = 0.32), we applied the PCA algorithm (MATLAB 2018v) to transform the original 10 local properties into lower dimensionality which could explain the major variation (above 90%) of the brain micrsotructure. The generated PCA maps were used to estimate the similarity between different vertices by calculating the Mahalanobis distance (Amunts et al., 2020; Zilles and Amunts, 2010) based on the PCA components, resulting in a 32k × 32k local-similarity matrix (32k standard meshes per hemisphere) for each hemisphere, namely PCA-based distance map. The spatial gradient was computed in workbench v1.5 for the local-similarity matrix, and the location with a high gradient represented the candidate areal borders. Then we used the “watershed by flooding” algorithm (Beucher and Lantuejoul, 1979) to identify the tentative boundaries in the gradient maps. The resulting 32k × 32k boundary map was further averaged over the second dimension to generate a border density map, which could quantify the probability of each spatial location being identified as a boundary (Gordon et al., 2016).

**Fig 1.**
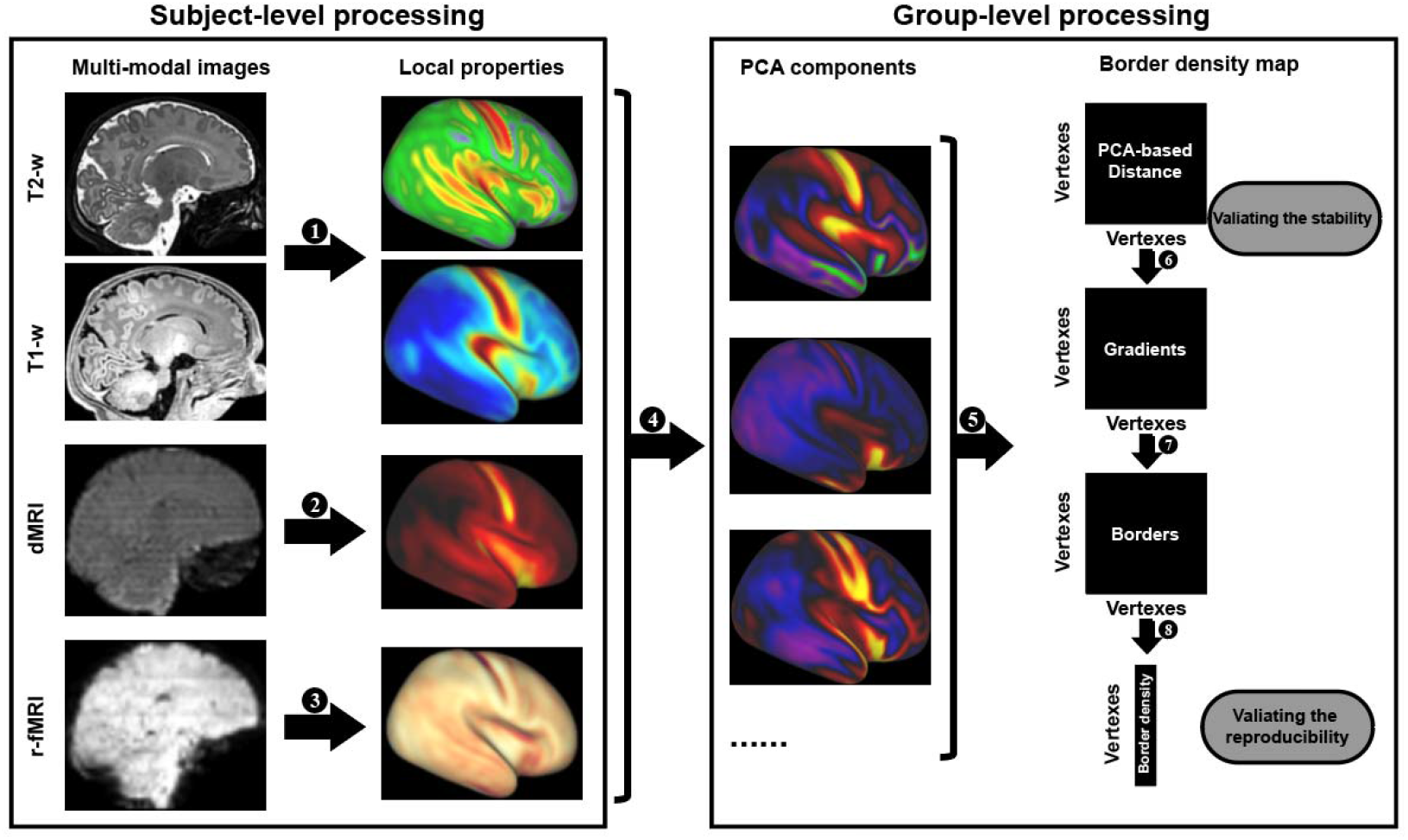
The flow chart of generating the multi-modal border density map. Acquiring 1) the cortical thickness and myelination from T1-w and T2-w data. 2) dMRI-derived properties, and 3) ALFF and fALFF from r-fMRI data. 4) Using PCA to combine different maps. 5) Calculating the distance map between vertices based on PCA-components. 6) Calculating the gradient maps from the distance maps. 7) Calculating the border density maps from the gradient maps. 8) Averaging the border density maps.

The border density map was the main reference in the next procedures generating the cortical parcellations, so we performed two necessary validation analyses before the boundary identification to assure the stability and reproducibility of the results.

### Stability of the boundary map

To test how the choice of 10 MRI features affects the final analysis (aka, stability of the parcellation), we estimated the contribution of each feature on the local similarity map by a leave-one-out approach. Specifically, we repeated the above procedures to generate a PCA-based distance map after we left one feature out, and then calculated the Pearson correlation between this new distance map and the original map generated by all features, to evaluate the influence of each feature on the final border density map. Meantime, we added two common properties of cortical morphology (e.g. the sulcus and curvature), and calculate the Pearson correlation between the two maps before and after adding the new features. In addition, we estimated the stability of the distance map by removing all derived measurements from a single modality, namely, CT and CM from structural MRI; ALFF and fALFF from r-fMRI; or tensor and NODDI parameters from dMRI, respectively. Finally, we directly measured the correlation between the original distance map and the map calculated through a single modality to estimate the contribution of a single modality on the distance map.

### Reproducibility of the boundary map

To determine if our parcellation was reliable at the group level, we separated the neonates into two groups with an equal number of subjects and repeated the above procedure to generate two border density maps, respectively. We then calculated the spatial correlation between the two groups in terms of the 10 MRI features and the derivate maps (e.g. averaged distance maps, averaged gradient maps, and border density maps). Finally, we generated the parcellation with the same parameters using the automated algorithm for the two groups and evaluated their agreement.

Particularly, since tiny differences in the border density maps could make a big difference in the final parcellations due to the sensitivity of the “automated algorithm”, the resulting labels and numbers of the parcellations would be different between the two groups, making it not feasible to calculate dice coefficient between matched regions in two groups. Therefore, for each ROI created in the group with less number of parcellation, we matched it to an ROI in the other group with the highest number of overlapped vertices, and calculated the dice coefficient between the matched ROIs between the two parcellations.

Given the quick development of the neonatal brain, we further estimated the effect of age on the results. Specifically, we separated the neonates into three groups according to their PMA at the scan and calculated the paired correlation coefficients of the averaged distance map among the three groups.

### Automated parcellation based on multimodal similarity

Both automated segmentation and manual delineation were used to generate the parcellations of the neonatal cortex. According to (Gordon et al., 2016), we generated the segmentation by identified all local minima on the bounder density map and then created the initial parcellation using a “watershed by flooding” procedure. The initial parcellation was further optimized by controlling the thresholds of the border density of the tentative borders and ensuring a minimum number of vertices (n=30) in a single parcel. Parcellations with different resolutions (300, 400, and 500 parcels) were generated by adjusting the threshold value of border density to adapt to different applications (Schaefer et al., 2018).

### Manual parcellation based on multimodal similarity

The manual method relied on the visually observable spatially continuous lines with relatively higher density values in the border density map. The boundaries were identified manually using the connectome workbench (v1.5) to obtain a parcellation with sufficient interpretability by a neuroanatomy expert (L.M.) and checked by another neuroanatomist (K.O.). The boundaries were drawn based on the following criteria.

1. All the boundaries should approximatively align to the border lines based on the border density map.
2. A tentative boundary should be in a similar location in both hemispheres to maximize symmetry in the parcellation.
3. For each tentative boundary, we would test whether the adjacent areas were statistically different from each other in the PCA-based distance. Particularly, we performed a two-sample t-test between the intra-parcel distance in two areas and the inter-parcel distance followed by a conservative Bonferroni corrected *p* < 0.05. The tentative boundary would be removed if the test was negative.
4. Furthermore, we also tested whether the adjacent parcels differed significantly in the original local measurements by a paired t-test across all subjects. Similarly, the boundary would be removed if none of the original properties could be distinguished between adjacent parcels after the Bonferroni correction.

The automated and manual parcellations at different resolutions are freely available on https://github.com/MingyangLeee/Neonatal-multimodal-parcellation.

## Results

### Stability of the PCA-based distance map

The population-averaged surface maps of 10 MRI features from 348 term-born neonates (PMA range from 37 to 41, mean age 39.93 ± 1.25; 164 females) showed distinct spatial patterns (Fig 1 and Supplementary Fig 1). We selected the first 5 components from PCA analysis of the original features since those components could explain more than 90% (94.02%, eigenvalues > 0.6) variation of the original data (all five PC maps were provided in the supplementary Fig 2). Prominent peaks of the first component were observed in the primary sensorimotor areas (motor, somatosensory, visual, and auditory areas), and the other peaks were located in the anterior and posterior cingulate gyrus and the precuneus. The second component had one peak in the parieto-occipital areas, especially in the occipital pole, and the other peaks in the crowns of the pre- and post-central gyrus. These PC maps were used to calculate the paired distance of vertices in the right and left hemispheres, respectively (Fig 2a). The results showed that the vertices with low distance to a particular location were not distributed isotopically but constrained in a specific anatomical structure (see supplementary Fig 3 for the distance map from sampled vertices).

**Fig 2.**
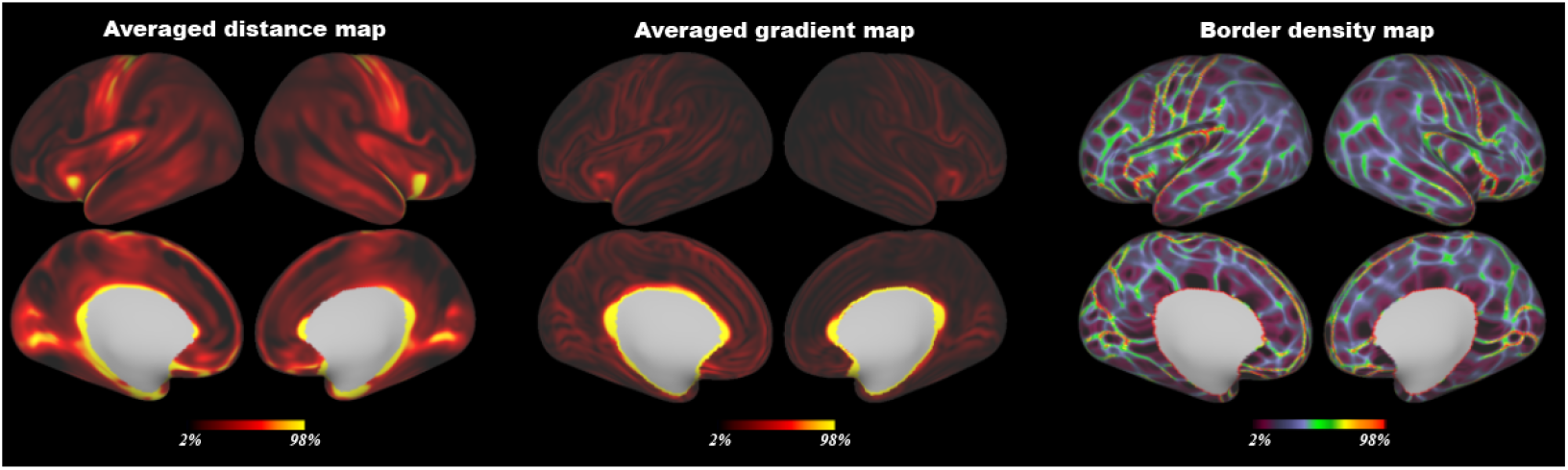
The averaged PCA-based distance map, the averaged gradient map, and the final border density map from all neonatal data.

To evaluate the stability of the distance map, we performed the leave-one-feature-out test. The spatial correlations between the leave-one-feature-out maps and the all-feature map were high in all the cases (*r* = 0.93 – 0.99, median value = 0.99, Table 2). We also tested the stability of the distance map by removing all derivate measurements from one MRI modality. The correlations were still high (*r* = 0.94 – 0.99, Table 2), except for the results of removing all dMRI-based properties (*r* = 0.81). Moreover, we measured the correlation between the distance map from all features and the map calculated through a single modality. The correlation was the highest in the tensor-based features (*r* = 0.82) and the lowest in the r-fMRI-based modality (*r* = 0.63; Table 2), suggesting the important contribution of dMRI data in the present parcellation. In addition, adding two common properties of cortical morphology (e.g. the sulcus and curvature) into the multimodal features did not change the similarity pattern (*r* = 0.973). Overall, those results suggested that the PCA-based distance map was relatively stable regardless of the choice of MRI features.

**Table 2.** The stability of the PCA-based distance map based on the correlations of the leave-one-out distance maps and the full-feature distance map.

| Property | Leave one property out | Modality | Leave one modality out | Correlation with one modality |
| --- | --- | --- | --- | --- |
| CT | 0.943 | Structure | 0.937 | 0.732 |
| CM | 0.986 |  |  |  |
| AD | 0.997 | Tensor | 0.939 | 0.817 |
| FA | 0.985 |  |  |  |
| MD | 0.998 |  |  |  |
| RD | 0.999 |  |  |  |
| ICV | 0.999 | NODDI | 0.986 | 0.783 |
| ODI | 0.988 |  |  |  |
| ALFF | 0.933 | r-fMRI | 0.946 | 0.627 |
| fALFF | 0.985 |  |  |  |

Abbreviations: CT = cortical thickness; CM = T1w/T2w-based cortical myelination; AD = axial diffusivity; FA = fractional anisotropy; MD = mean diffusivity; RD = radial diffusivity; ICV = intra-cellular volume fraction; ODI = orientation dispersion; ALFF = amplitude of low-frequency fluctuations; fALFF = fractional ALFF.

### Reproducibility of the border density map

To determine the reliability of the bounder maps, we separate the datasets into two equal groups (*n* = 174 for each group; PMA at scan = 41.06 ± 1.80 and 40.85 ± 1.61; *t* = 1.15, *p* > 0.2). The two groups showed high consistency in all 10 original properties (*rs* = 0.991 – 0.999, Table 3), the averaged distance map (*r* = 0.995), the averaged gradient map (*r* = 0.999), and the border density map (*r* = 0.943), indicating high reliability of the boundary information used for generating the parcellation. Finally, we generate the parcellations using the two border density maps from two groups. The first parcellation contained 269 regions (right: 122, left: 147) while the second parcellation resulted in 273 regions (right: 124, left: 149). The proportion of overlapped vertices between matched ROIs in two groups was 0.79. Of the two parcellations, 232 regions could be matched, and the dice coefficient of the machted ROIs were 0.77 ± 0.20.

**Table 3.** The reproducibility of the surface maps between two neonatal groups.

| **Cortical property** | **correlation between two groups** |
| --- | --- |
| **CT** | **0.9957** |
| **CM** | **0.9981** |
| **AD** | **0.9957** |
| **FA** | **0.9921** |
| **MD** | **0.9958** |
| **RD** | **0.9956** |
| **ICV** | **0.9941** |
| **ODI** | **0.9913** |
| **ALFF** | **0.9959** |
| **fALFF** | **0.9923** |
| **Distance** | **0.9951** |
| **Gradient** | **0.9986** |
| **Border density** | **0.9425** |

Abbreviations: CT = cortical thickness; CM = T1w/T2w-based cortical myelination; AD = axial diffusivity; FA = fractional anisotropy; MD = mean diffusivity; RD = radial diffusivity; ICV = intra-cellular volume fraction; ODI = orientation dispersion; ALFF = amplitude of low-frequency fluctuations; fALFF = fractional ALFF.

We further tested the effect of age on the density map by separating the subjects into three groups according to their PMA at scan (*n* = 116 in each group, mean PMA = 43.04, 41.11, and 39.22, respectively), and the paired correlation coefficients of the averaged distance map among the three groups were high (0.95 – 0.98). This result suggested that the proposed PCA-based distance map was relatively stable across the neonatal period, so we could use the averaged data from all neonates.

### Automated parcellation of the neonatal cortex at multiple resolutions

By adjusting the thresholds of border density based on the gradient of the distance map of multi-modal MRI properties, the proposed algorithm could generate the parcellations with a different number of parcels. Three typical resolutions were present in Fig 3 and Supplementary Fig S4, which resulted in 500 (right: 233; left: 267), 400 (right: 186; left: 214), and 300 (right: 133; left: 167) parcels, respectively.

**Fig 3.**
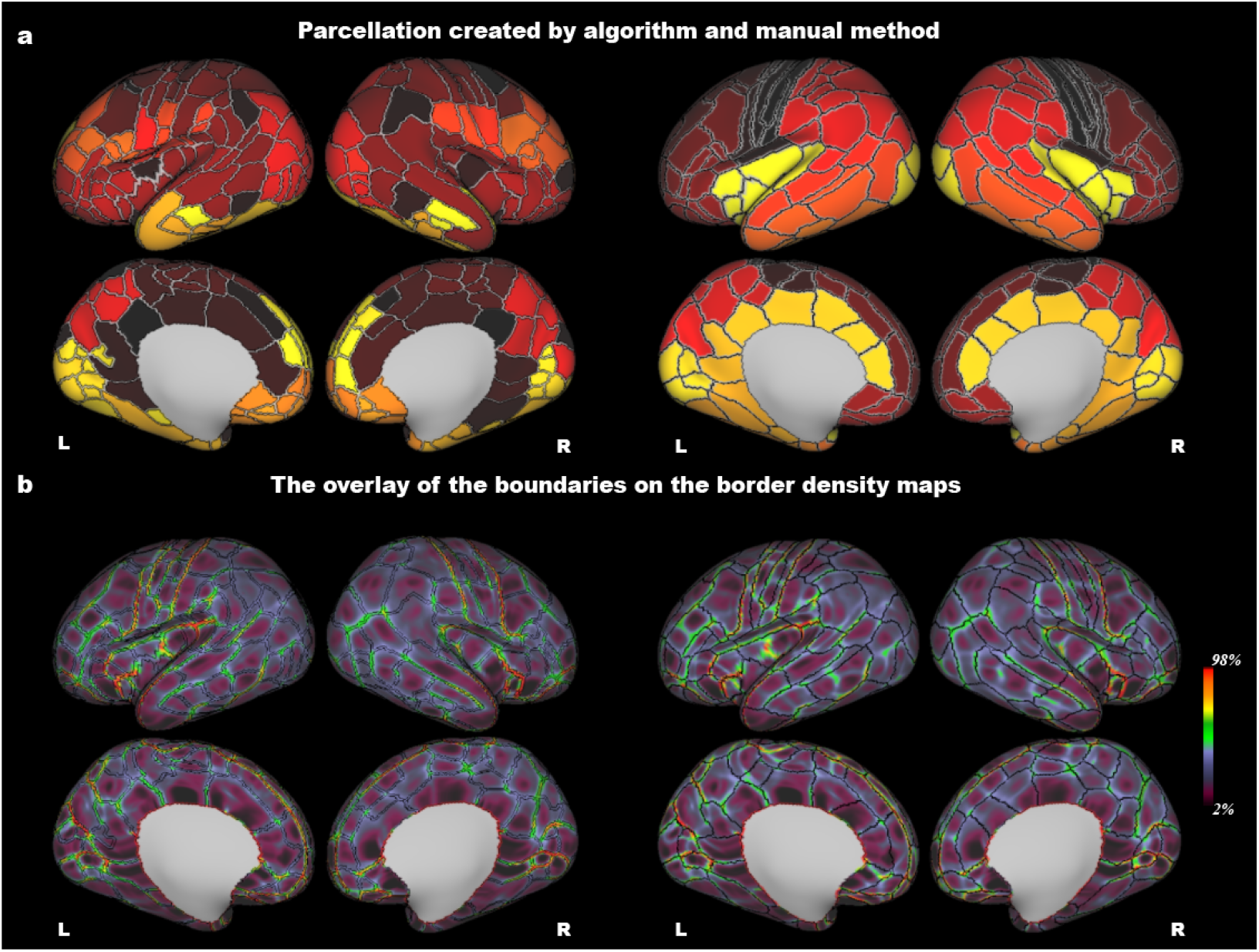
a) The parcellations which were created by an automatic algorithm (300 parcels) and the manual method. b) The overlay of the boundaries on the border density maps.

### Neuroanatomy-based manual parcellation of the neonatal cortex

The manual parcellation resulted in 105 cortical areas in each hemisphere (Fig 3 and figure S4-5), respectively. We provided a detailed description of the anatomical location in Supplementary table 1. Briefly, the number of vertices in each parcel was 204±125, ranging from 25 to 650 vertices. The correlation between the number of vertexes and the intra-parcel distance of the parcels was not significant (*r* = 0.05 and 0.08; *p* > 0.35 in the right and left hemispheres, respectively), which indicated that thelocal similarity was not significantly influenced by the size of parcels. For any of the adjacent parcel pairs, the intra-parcel distance was significantly lower than the inter-parcel distance (*p*-values < 10^−6^ after Bonferroni correction; Fig 4a), and at least one of the 10 original features showing significant differences between the adjacent areas (Bonferroni corrected *p* < 0.05; Fig 4b). Furthermore, we performed a sanity check for the consistency between the borders and the gradients of used properties. As shown in figure S7-8, most boundaries of the final parcellation are located on the peaks of gradient maps of the original features.

**Fig 4.**
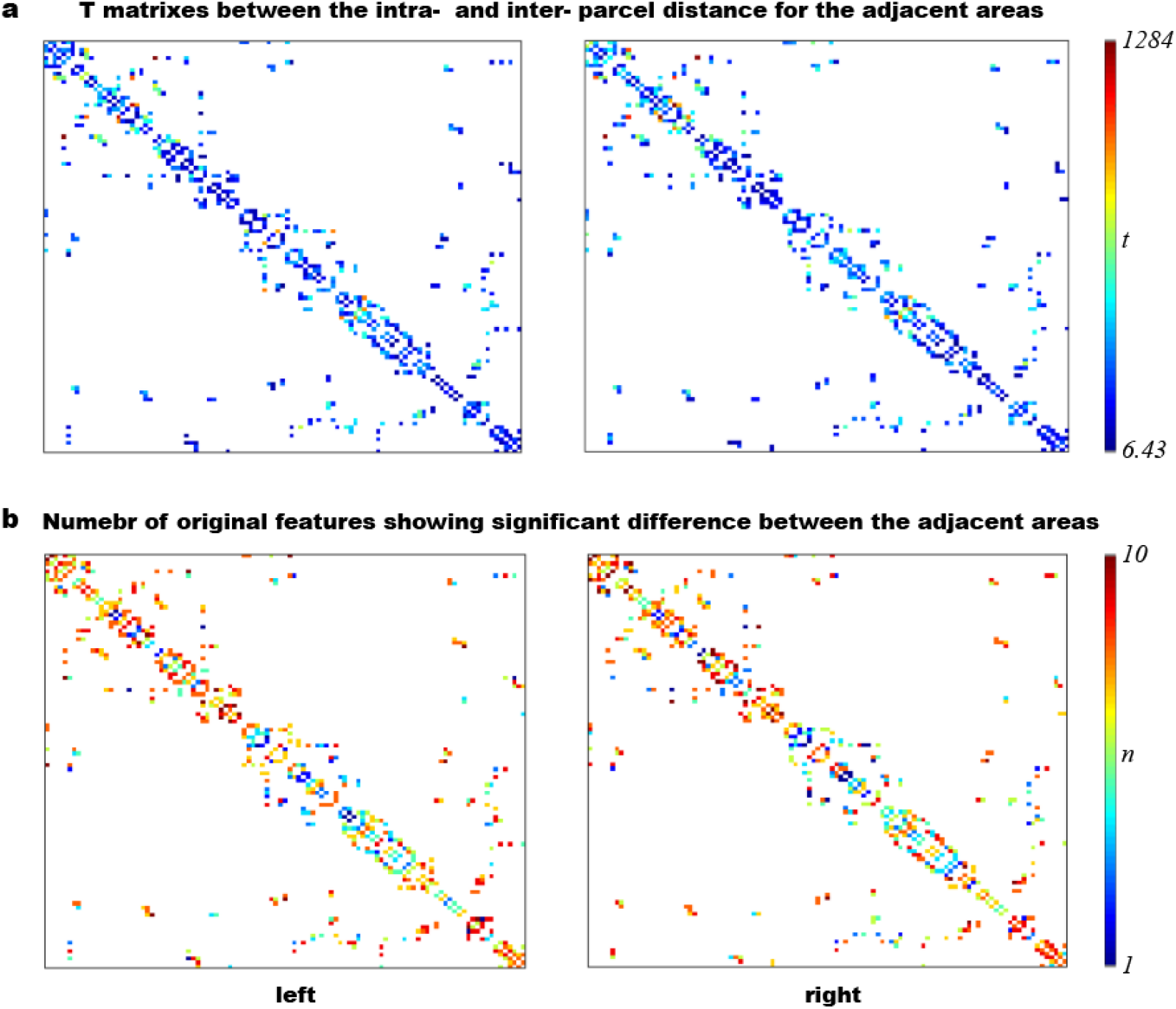
The statistical results of the manual parcellation based on difference between the adjacent parcels. The x and y axes represent 105 cortical regions and the labels of those regions are shown in supplementary table 1 in order. The non-empty (colored) entry [i, j] in the matrix indicates that the area i and j are adjacent. The value in the matrix denotes the t value of the difference between intra- and inner-parcel distance for the adjacent areas a), or the number of original features showing significant differences between the adjacent areas b).

## Discussion

In this study, we proposed a multi-modal similarity-based approach for parcellating the human neonatal cerebral cortex. The border information generated from the integrated MRI features was relatively stable with different MRI features and reproducible at the group level. Therefore, we could use the border information to generate multi-modal multi-resolution parcellations for future studies with different research purposes. We performed an automatic analysis to create three parcellations with 500, 400, and 300 parcels, and also manually create a parcellation with high anatomical interpretability (210 parcels). The proposed parcellations have more spatial details compared to the previous parcellation based on microanatomy (Adamson et al., 2020; Alexander et al., 2017; Feng et al., 2019; Gousias et al., 2013, 2012; Shi et al., 2011) and were more interpretable compared to the parcellation based on functional connectivity (Shi et al., 2018; Wang et al., 2021). More importantly, the unbiased parcellation by integrating macro-structural (from T1w/T2w), functional (from r-fMRI), and micro-structural (from dMRI) features would provide an ideal reference atlas in the structural-functional analysis of the developing human connectome.

In our view, the fundamental difference among various brain parcellations mainly lies in the information used to generate the boundaries. One unique contribution of the current parcellation is that we used PCA-based distance information from multi-modal MRI images instead of single modality as used in the other neonatal atlases (Alexander et al., 2017; Feng et al., 2019; Gousias et al., 2012; Shi et al., 2011; Wang et al., 2021), and thus the generated atlas could be particularly useful for unbiased functional-structural analysis. Previous studies suggested that the cortical myelination obtained from the T1-weighted and T2-weighted contrast (Glasser and van Essen, 2011) was suggested to be correlated with the degree of myelination (Soun et al., 2017), which was already widely used in both adult and infant study (Caspers et al., 2013; Fenchel et al., 2020; Glasser et al., 2013; Makropoulos et al., 2018; Nazeri et al., 2022; Soun et al., 2017). dMRI reveals the tissue microstructure based on restricted water diffusion (Mori and Zhang, 2006). For example, the NODDI model (Zhang et al., 2012) could quantitatively estimate the neurite density, orientation dispersion, and fractional CSF indexes, which may be related to the cytoarchitecture and myeloarchitecture (Fukutomi et al., 2018). The r-fMRI-based measurements measure the averaged amplitude of the BOLD signal, reflecting the microenvironment and overall metabolism of local tissues (Zou et al., 2008). Therefore, the integration of these features could better present the microstructural similarity of cortical vertices from a wider range of perspectives and improve the stability of the boundary estimation.

Using PCA, we kept the top components that could explain more than 90% variation across the original features. Those components could provide the spatial information that was shared among the multiple imaging modalities and thus were relatively stable. This PCA-based distance used here was analogical to the morphometric similarity network (Fenchel et al., 2020; Seidlitz et al., 2018), which was used to estimate the covariate of various structural properties such as sulcus, cortical thickness, and cortical myelination, while here we added diffusion and functional features to enable cross-modal analysis. As shown in Figure 2, the areas with maximum averaged distance across the whole cortex were located in the primary sensorimotor system such as motor, somatosensory, visual, and auditory areas, while the areas with minimum averaged distance were sparsely distributed in the frontal (e.g. inferior frontal gyrus and middle frontal gyrus) and parietal lobe (e.g. precuneus and inferior parietal lobe). This pattern was similar to previous studies characterizing the sensorimotor-to-association axis of the cortical orgnization (Margulies et al., 2016; Sydnor et al., 2021; Vázquez-Rodríguez et al., 2019), which showed a principal gradient from the transmodal regions (the default-modal network) to the unimodal areas (sensorimotor areas) in both human and monkey brain. Together, those results suggested that different measurements from various MRI modalities may share common patterns (Kelly et al., 2012) for us to create the multi-modal parcellation in the present study.

There could be different methods for creating the final parcellation with the PCA-based distance map. In this study, we mainly followed Gordon et al. (Gordon et al., 2016) for automated parcellation, which used a “watershed by flooding” algorithm to detect the local ridge of the gradient map. This method has been successfully applied in previous studies to generate functional connectivity based parcellations in human adults. (Cohen et al., 2008; Gordon et al., 2016; Johnson et al., 2009) and the resulting parcellations showed relatively good homogeneity (Gordon et al., 2016; Schaefer et al., 2018). However, the data-driven parcellations were sensitive to the parameters applied in the algorithm, and easily produced parcels with low interpretability. So we additionally provided a manual delineated neonatal cortical parcellation. The manual method employed prior knowledge about neuroanatomy and we can filter out the asymmetrical boundaries and the boundaries driven by noise (Glasser et al., 2016), but the subjective factor during the delineation was the main consideration in the present study as there was no gold standard. Nevertheless, considering that the meaning of a boundary is to distinguish specific attributes, we applied strict quantitative criteria to make sure that each of the final boundaries could distinguish both the PCA-based distance and at least one of the original properties. Although this approach could not completely avoid the variation of the boundaries driven by subjective factors, it guaranteed that the final boundaries were relatively biophysically meaningful. Note that the present parcellation is largely different from the previous parcellations based on macroanatomy (e.g. MCRIB or AAL), which were defined based on the major sulcus of the cortex (Alexander et al., 2017; Desikan et al., 2006; Shi et al., 2011; Tzourio-Mazoyer et al., 2002). The PCA-based distance map showed that the vertices on the two slides of a sulcus may have similar MRI features, and thus the present method could result in the boundaries located between the sulcus and gyrus.

There are several limitations in the current work. First, the parcellation created in the current study shared a common pattern but also showed distinct differences from the previous single-modal-based parcellations. One typical difference could be observed in the visual cortex, which did not exactly follow the visuotopic definition from r-fMRI data (Arcaro and Livingstone, 2017; Glasser et al., 2016). We also found that the myelination map provided a good contrast for the border of the V1 area, but the integrated border density map did not. Future work could flexibly reweight the different modalities for a particular area to obtain a parcellation consistent with the classical or cytoarchitectural atlas. In addition, due to the lacking of task-fMRI data in the neonatal population, we were not sure whether the visuotopic mapping in human neonates was similar to human adults because the central fovea was not fully developed in the neonatal period (Candy et al., 1998). The second limitation came from the manual method which could not be easily applied to a new population or individual subjects. A previous adult brain study first used the manual method to obtain an initial parcellation and then trained a machine learning classifier to learn the critical features in the parcels (Glasser et al., 2016), which may be used to validate the generalizability of the manual parcellation. The third limitation lies in the image quality and registration process. Although dHCP provided a state-of-the-art pipeline for human neonates (Bozek et al., 2018; Makropoulos et al., 2018; Robinson et al., 2018), the quality of the original images from infants was not as good as the adults, due to neonatal motion, low contrasts, and insufficient resolution in neonatal brain MRI (Hughes et al., 2017). This inevitably influenced the segmentation and registration preprocessing steps. We directly use the transformation files from the preprocessing including a volume-surface transformation and an individual-template transformation in the surface space, which might accumulate inaccuracy in the final results. The present parcellation might be improved given higher quality data and advanced preprocessing algorithms.

## Conclusion

We proposed multi-modal-based cortical parcellations designed for the neonatal brain. The PCA-based distance map used for parcellation integrated 10 MRI features from at macrostructural, microstructural, and functional levels, which was shown to be stable to the choice of MRI features and repeatable to the neonatal populations. We utilized this cross-modal information to generate the neonate-specific parcellations by an automatic algorithm at multiple resolutions (300-500 parcels), as well as manually delineated parcellations with good interpretability and asymmetry, to be adaptable to various needs in future studies about the development of human connectome.

