## Supplemental figures and table for "Multi-modal multi-resolution atlas of the human neonatal cerebral cortex based on microstructural similarity"

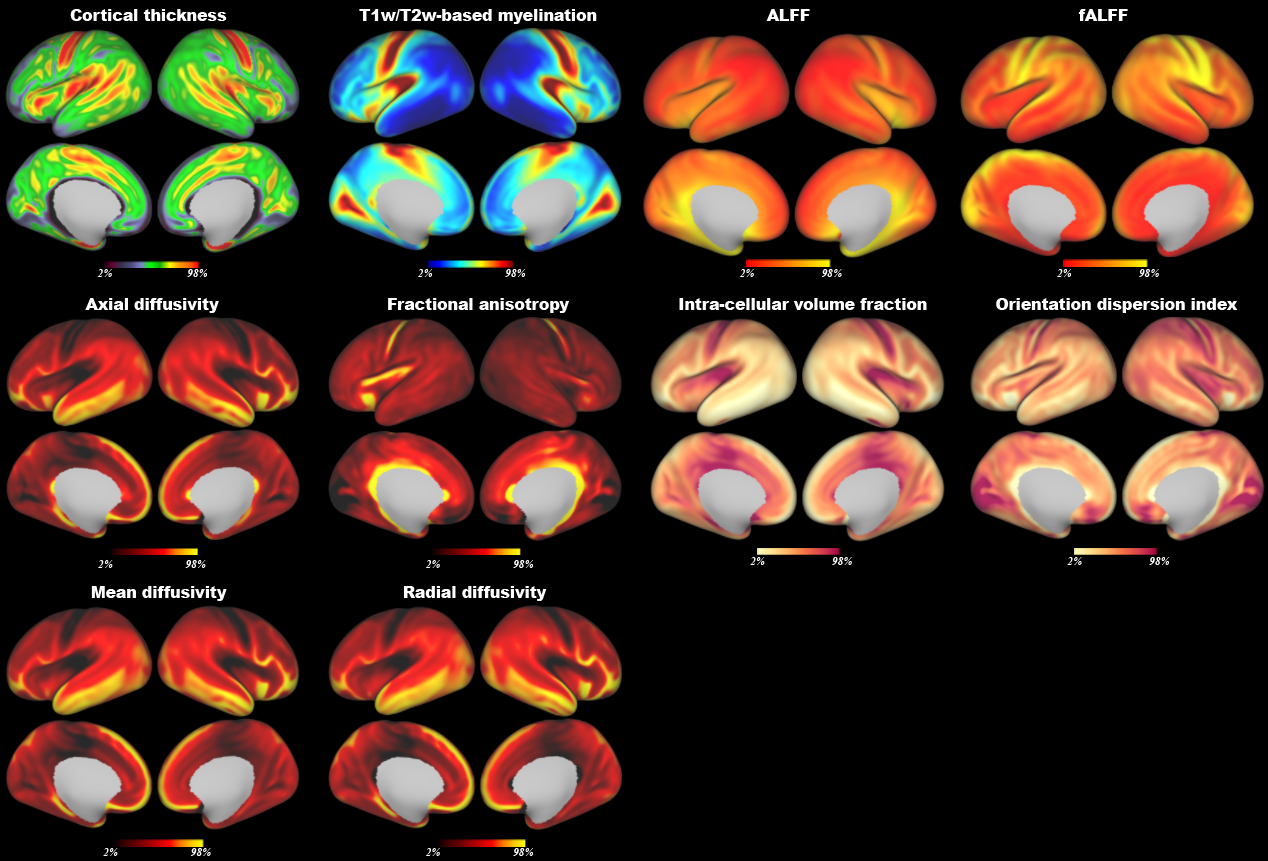

Fig S1. The multimodal MRI feature maps averaged across all neonates in the present study, including cortical thickness and T1w/T2w myelination from structural MRI, axial diffusivity, fractional anisotropy, mean diffusivity, radial diffusivity of the tensor model and intra-cellular volume fraction, orientation dispersion of the NODDI model from diffusion MRI; and amplitude of low-frequency fluctuations (ALFF), fractional ALFF from rest-state functional MRI.

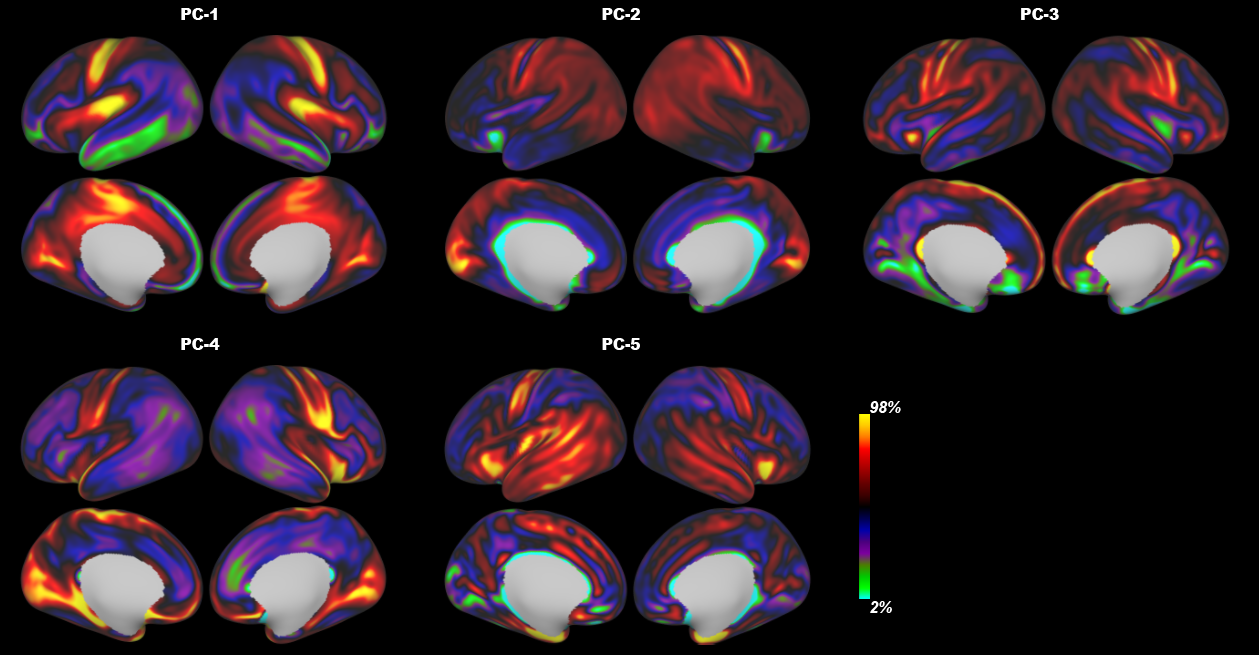

Fig S2. The top five principal component maps were obtained from the 10 MRI features from three MRI modalities. Those components could explain more than 90% (94.02%, eigenvalues > 0.6) variation of the original data.

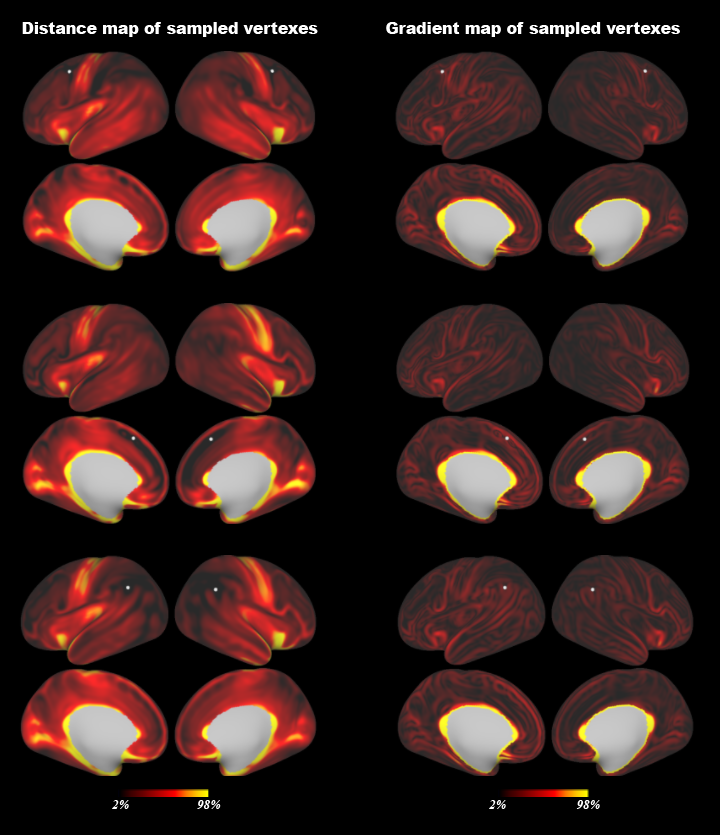

Fig S3. Sampled distance maps of three exemplary vertices (White dots). The gray sphere indicates the source vertex and the color describe the PCA-based distance between the source vertex and other vertices (e.g. darker colors indicate a closer distance).

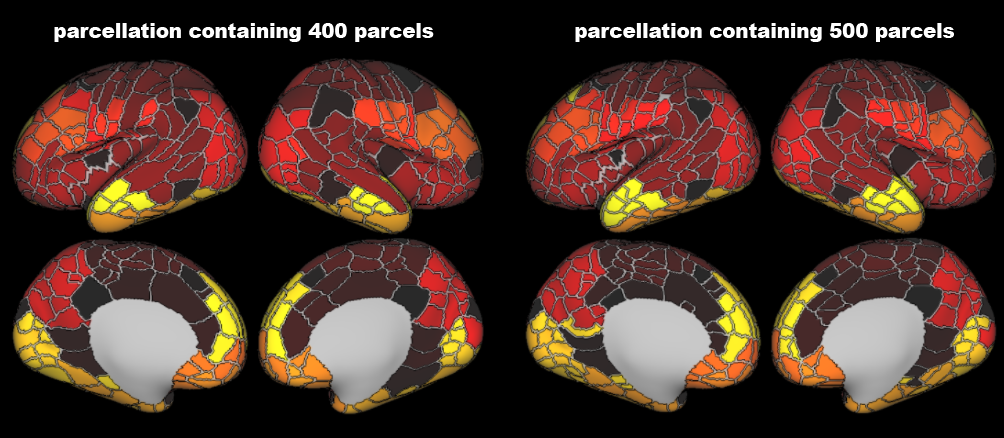

Fig S4. The parcellations with 400 (left) and 500 (right) parcels are created by an automatic algorithm.

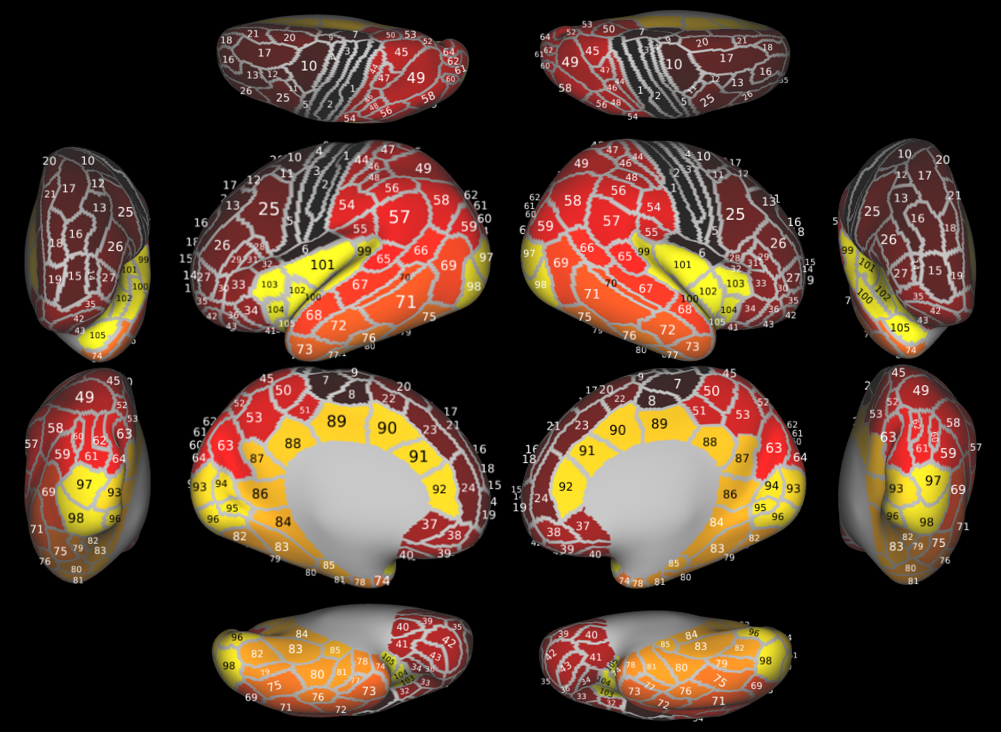

Fig S5. The manual parcellations in multiple views on the flat surface, the labels of the areas are shown in supplementary Table 1.

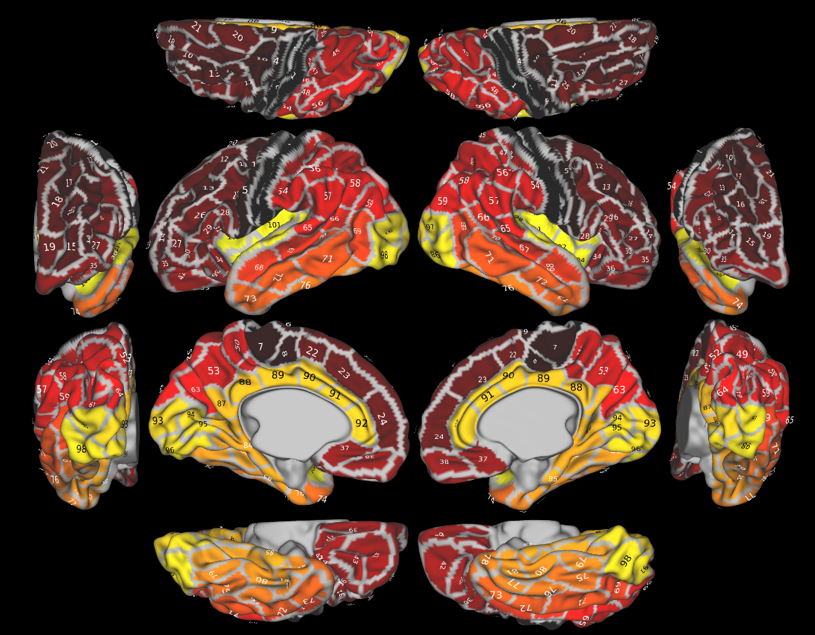

Fig S6. The manual parcellations in multiple views on the original surface, the labels of the areas are shown in supplementary Table 1.

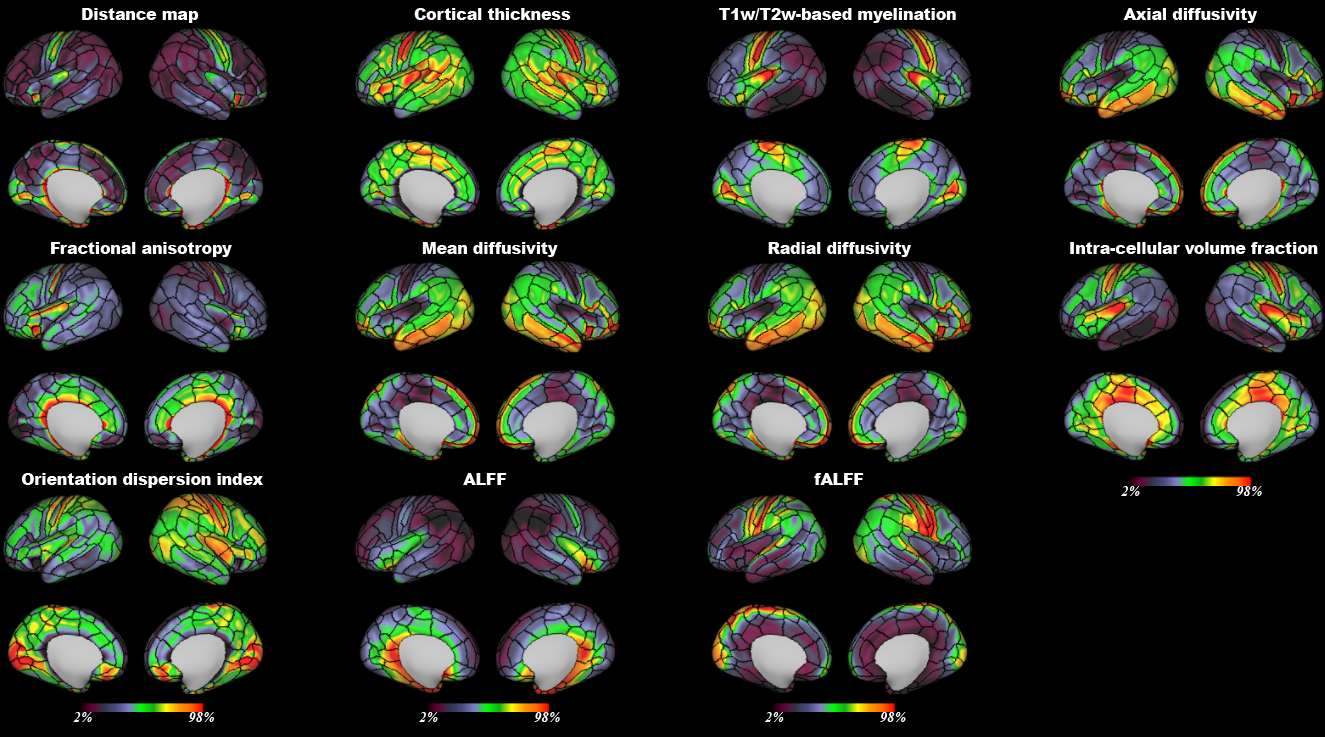

Fig S7. Overlap of manually drawn boundaries on the individual feature maps averaged across all neonates.

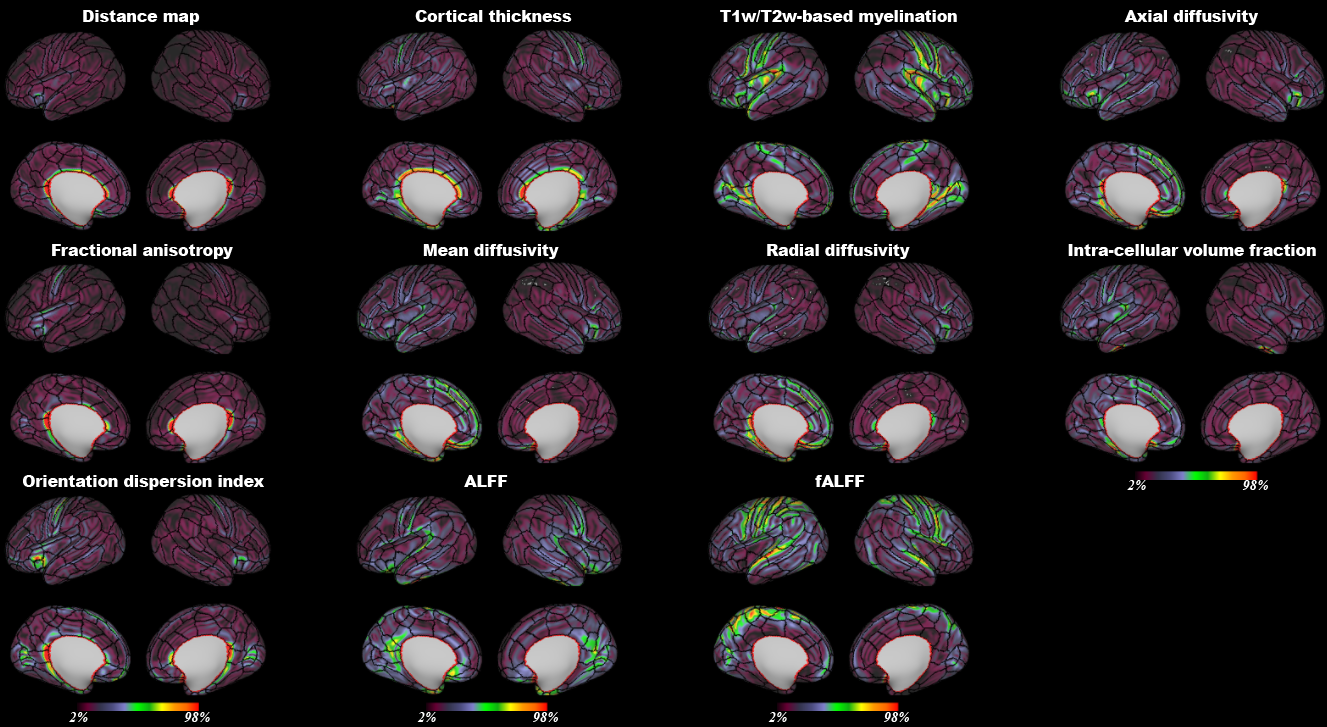

Fig S8. Overlap between manually drawn boundaries on the gradient maps of individual MRI features.

Table S1. The lists of the 105 regions of the manual parcellation.

| abbreviation | description | Index |
| --- | --- | --- |
| PoCG1 | posterior part of the postcentral gyrus | 1 |
| PoCG2 | anterior part of the postcentral gyrus | 2 |
| PrCG1 | posterior part of the precentral gyrus | 3 |
| PrCG2 | anterior-superior part of the precentral gyrus | 4 |
| PrCG3 | anterior-inferior part of the precentral gyrus | 5 |
| OP_c | the opercular area inferior to the central region | 6 |
| PCL1 | posterior part of the paracentral lobule | 7 |
| PCL2 | anterior part of the paracentral lobule | 8 |
| SF_sp | superior-posterior part of the superior frontal gyrus | 9 |
| PRS_s | superior part of precentral sulcus | 10 |
| MFG1 | four parts of the middle frontal gyrus from posterior to anterior segments | 11 |
| MFG2 |  | 12 |
| MFG3 |  | 13 |
| MFG4 |  | 14 |
| IMFS_v | vertical segment of the intermediate frontal sulcus | 15 |
| IMFS_h | horizontal segment of the intermediate frontal sulcus | 16 |
| SFS1 | posterior part of the superior frontal sulcus | 17 |
| SFS2 | anterior part of the superior frontal sulcus | 18 |
| SFG_s1 | anterior part of the superior frontal gyrus | 19 |
| SFG_s2 | posterior part of the superior frontal gyrus | 20 |
| SFG_s3 | middle part of the superior frontal gyrus | 21 |
| SFG_m1 | three part of the medial superior frontal gyrus from posterior to anterior segments | 22 |
| SFG_m2 |  | 23 |
| SFG_m3 |  | 24 |
| PRS_i | infeiror part of the precentral sulcus | 25 |
| IFS1 | posterior part of the inferior frontal sulcus | 26 |
| IFS2 | anterior part of the inferior frontal sulcus | 27 |
| IFG1 | three part of the inferior frontal gyrus from posterior to anterior segments | 28 |
| IFG2 |  | 29 |
| IFG3 |  | 30 |
| FOP1 | posterior-medial part of the frontal opercular area | 31 |
| FOP2 | posterior-lateral part of the frontal opercular area | 32 |
| FOP3 | middle part of the frontal opercular area | 33 |
| FOP4 | anterior part of the frontal opercular area | 34 |
| LORG | lateral orbital gyrus, the majority of this area is in the inferior frontal gyrus | 35 |
| IFG_or | orbital part of the inferior frontal gyrus | 36 |
| PTG | paraterminal gyrus | 37 |
| ROG | rostral gyrus | 38 |
| GR | gyrus rectus | 39 |
| OLF | olfactory sulcus | 40 |
| PORG1 | three part of the orbitofrontal gyrus | 41 |
| PORG2 |  | 42 |
| PORG3 |  | 43 |
| PCS1 | superior part of the postcentral sulcus | 44 |
| SP1 | superior part of the superior parietal area | 45 |
| PCS2 | middle-posterior part of the postcentral sulcus | 46 |
| SP2 | inferior part of the superior parietal area | 47 |
| IPS1 | anterior-inferior part of the inferior parietal sulcus | 48 |
| IPS2 | main part of the inferior parietal sulcus | 49 |
| PreCu1 | anterior-superior part of the precuneus | 50 |
| PreCu2 | anterior-infeior part of the precuneus | 51 |
| SP3 | superior-posterior part of the superior parietal area | 52 |
| PreCu3 | posterior part of the precuneus | 53 |
| PCS3 | inferior part of the postcentral sulcus | 54 |
| OP_p | posterior part of the opercular area | 55 |
| SMG | superior part of the supramarginal gyrus | 56 |
| TPO | temporal-parietal-occipital junction | 57 |
| AG | Angular gyrus | 58 |
| IPL | posterior-inferior part of the inferior parietal lobe | 59 |
| IPS3 | three part of the posterior inferior parietal sulcus from lateral to medial segments | 60 |
| IPS4 |  | 61 |
| IPS5 |  | 62 |
| POS | main part of the parieto-occipital sulcus | 63 |
| SOG | Superior occipital gyrus | 64 |
| STG_p1 | two part of the posterior superior temporal gyrus | 65 |
| STG_p2 |  | 66 |
| STG_m | middle part of the superior temporal gyrus | 67 |
| STG_a | anterior part of the superior temporal gyrus | 68 |
| AOS | anterior occipital sulcus | 69 |
| STS_i | infeiror part of the superior temporal sulcus | 70 |
| TE_p | posterior temporal lobe | 71 |
| MTG | middle temporal gyrus | 72 |
| TP_dl | dorsal-lateral part of the temporal pole | 73 |
| TP_dm | dorsal-medial part of the temporal pole | 74 |
| OTS_p | posterior part of the occipitotemporal sulcus | 75 |
| ITG_p | posterior part of the inferior temporal gyrus | 76 |
| ITG_a | anterior part of the inferior temporal gyrus | 77 |
| TP_v | ventral part of the temporal pole | 78 |
| FUS_p | posterior part of the fusiform gyrus | 79 |
| FUS_m | middle part of the fusiform gyrus | 80 |
| FUS_a | anterior part of the fusiform gyrus | 81 |
| COS_p | posterior part of the collateral sulcus | 82 |
| COS | main part of the collateral sulcus | 83 |
| PHA_p | posterior part of the parahippocampal gyrus | 84 |
| PHA_a | anterior part of the parahippocampual gyrus | 85 |
| Cing1 | seven part of the cingulate gyrus from posterior to anterior segments | 86 |
| Cing2 |  | 87 |
| Cing3 |  | 88 |
| Cing4 |  | 89 |
| Cing5 |  | 90 |
| Cing6 |  | 91 |
| Cing7 |  | 92 |
| cuneus | main part of the calcarine sulcus | 93 |
| calcarine | anterior-superior part of the calcarine sulcus | 94 |
| lingual1 | anterior part of the lingual gyrus | 95 |
| lingual2 | posterior part of the lingual gyrus | 96 |
| MOG | middle occipital gyrus | 97 |
| IOG | inferior occipital gyrus | 98 |
| TPM | the planum temporale | 99 |
| TTG | Transverse temporal gyrus | 100 |
| INS_ps | superior part of the posterior insula | 101 |
| INS_pi | inferior part of the posterior insula | 102 |
| INS_as | Superior part of the anterior insula | 103 |
| INS_am | Middle part of the anterior insula | 104 |
| INS_ai | Inferior part of the anterior insula | 105 |
